# Spatial and Temporal Patterns of Wolf [*Mahihkan* (Cree), *Tha* (Denesuline), *Amaruk* (Inuktitut), *Canis lupus*] Occurrences on the Summer Range of the Eastern Migratory Cape Churchill Caribou Population in the Hudson Bay Lowlands of Manitoba

**DOI:** 10.1101/2025.04.11.648486

**Authors:** R.K. Brook, K.E Harris, D.A. Clark, C. Lochansky, J.A. Colpitts

**Affiliations:** Department of Animal and Poultry Science, College of Agriculture and Bioresources, University of Saskatchewan, 51 Campus Drive, Saskatoon, Saskatchewan, S7N 5A8, Canada; School of Environment and Sustainability, University of Saskatchewan, 117 Science Place, Saskatoon, Saskatchewan S7N 5C8

**Keywords:** *Canis lupus hudsonicus*, Eastern Migratory caribou, Hudson Bay, non-invasive, predators, *Rangifer tarandus*, sub-arctic, trail cameras, Wapusk National Park, wolves

## Abstract

Wolves (*Canis lupus*) function as a top predator across diverse ecosystems including the sub-arctic, and they have been managed in often controversial ways. Communities and scientists are increasingly supporting minimally invasive research and monitoring, including using trail cameras. We employed a network of 15 Reconyx trail cameras at three monitoring areas aimed at detecting the spatial and temporal aspects of wolf occurrences within the summer range of Eastern Migratory Cape Churchill caribou in Wapusk National Park in the Hudson Bay Lowlands of Manitoba, Canada from 2013-2021. In this first peer-reviewed quantitative study of wolves in the region, we found that wolves detection events were generally consistent across years. Wolf distribution was consistently positively skewed toward the southern part of the caribou summer range in all years. Wolves experienced extreme environmental conditions, with a 60°C range in temperature, from a low of −32°C in winter to a high of +28°C in summer and an annual change in day length of >11 hours between summer and winter. Wolves occurred most commonly in spring and summer and occurred at equal frequency during night and day overall but selected for nighttime in September, October, and November as day length shortened dramatically.

## Introduction

Wolves (*Canis lupus*), referred to in northern Canada by Indigenous communities as *Mahihkan* by Cree People, *Tha* by Denesuline People, and *Amaruk* by Inuktitut People, are a keystone ecological and cultural species across their global range, having top-down effects on entire ecosystems (Schmitz et al. 2010, Winnie and Creel 2017, Anderson et al. 2024) and influencing species such as caribou (*Rangifer tarandus*; Dalerum et al. 2017; Mishra 2024). Management of wolves across North America has been highly controversial over the last century and beyond, with past and present efforts often focused on over-simplified mechanistic approaches that dubiously presumed purposely killing wolves would result in increased populations of ungulates, particularly caribou (*Manitoba Municipal Act,* 1873; Johnson et al. 2022). Across their North American range, wolves have been consistently blamed for livestock depredation and declines in free-ranging ungulate populations, often without evidence, and killed in large numbers legally and illegally regardless of reality (Barclay 2010, Proulx and Rodtka 2015, Proulx and Parr 2018, Blossey and Hare 2022). This approach has been particularly contentious over the last century due to the use of methods such as strychnine poison, trapping, and shooting from helicopters that challenge scientists’ and the public’s notions of ethics and animal welfare (Brook et al. 2015).

Contemporary wolf research and management has been challenged to adopt less invasive approaches, particularly in northern Canada where Indigenous people have raised significant concerns about conventional wildlife research and management approaches such as collaring, aerial surveys, and population models (Wolfe 2006, Freeman and Foote 2009). Trail cameras have been identified by scientists and communities as a non-invasive (Barber-Meyer 2022) or minimally invasive tool to conduct long-term research on wildlife, including wolves (Hansen and Urbigkit 2021; Ausband et al. 2022; Clare et al 2023). Trail cameras can function continuously, collecting observations across seasons and among years (Clark et al. 2019). Trail cameras also require minimal fieldwork; they are cost effective; they collect data passively while animals move past them; and they can be visited as little as once per year (Laforge et al. 2017, Lochansky et al. 2025).

Understanding wolf occurrences, distribution, and movements is critical for understanding caribou ecology as wolves have complex interactions with caribou across spatial scales that include direct predation as well as indirect effects, including initiating trophic cascades (Ripple et al. 2025). The overall wolf range in North America overlaps completely with the distribution of migratory caribou, and wolves are the most common predator of caribou (Muisani et al. 2007; Michelot et al. 2023; Government of Canada 2024). Wolves in northern Canada are characterized by large scale movements, with Walton et al. (2001) finding satellite collared wolves to have an average home range size of 63,000 km^2^. Latham et al. (2011) determined that wolves follow natural linear features, including streams and rivers, as these areas provide ease of movement and abundant prey. In the sub-arctic, wolves are typically not territorial and move with migratory caribou from summer tundra to winter forest-tundra (Muisani et al. 2007).

Wolves may make extremely large foraging movements; Frame et al. (2004) documented a Global Positioning System (GPS) collared adult female wolf in the Northwest Territories making a 341 km foray over two weeks from the tundra denning area to an area with Bathurst caribou and back again. While it is well-established that wolves prey upon caribou, sometimes travelling long distances to do so, long-term spatial and temporal patterns of wolf occurrences remain generally poorly understood.

Wolf activities vary seasonally and in response to numerous environmental variables. Bi-modal crepuscular activity peaks have been documented in Alaska (Fancy and Ballard 1995) and Poland (Theuerkauf et al. 2003). In contrast, wolves have been documented being active throughout the night (Chavez and Gese 2006) or even exhibiting daytime forays (Vilà et al. 1995). These differences in peaks of activity have been linked to human presence, representing a threat to predators (Chavez and Gese 2006; Botts et al. 2020), and latitude (Vilà et al. 1995; Packard 2006; Rossa et al. 2021), dictating daylight hours and annual mean temperature. During the twilight hours, the intensity of moonlight shifts with the lunar phases, shaping low light hunting strategies by affecting prey sightings, predator success, and prey vigilance (Boiseau et al. 2024). As a result of the wide range of factors influencing wolf movements on a daily, seasonal, annual, and multi-annual basis, specific details of wolf activity should be directly observed in order to determine patterns and responses to fluctuating periods of daylight and darkness.

During the winter months in high-latitude environments with severely limited daylight hours, such as the Arctic tundra, wolves may show less distinct crepuscular peaks compared to their lower-latitude counterparts (Vilà et al. 1995; Packard 2006; Rossa et al. 2021). Here, extended periods of twilight compress the window of true darkness, potentially leading to a broader activity window throughout these twilight hours, allowing wolves to maximize their hunting opportunities even when daylight hours are scarce. Other influences on wolf activity identified thus far include temperature, light intensity, and prey activity (Packard 2006). Especially at high latitudes like Alaska, activity patterns shift with the seasons, with stronger crepuscular patterns observed in summer (peaks at 0600 and 2200) than winter, possibly to avoid high temperatures, and lower overall activity in winter (Fancy and Ballard 1995). This adaptability underscores the ability of wolves to adjust their behaviour based on the specific ecological pressures they face. It is likely these patterns occur in wolves in northern Canada as well, but it is yet to be directly studied in Manitoba. Wolves are present across virtually all of Manitoba, on agriculture lands to the south, through the boreal forest and into the northern woodlands tundra, up to the Nunavut border. The northern half of Manitoba has widespread and overlapping wolf and caribou populations. Despite their importance as a keystone ecological species and keystone cultural species, there has been very little research done on wolves in northern Manitoba. Indeed, there is only one existing peer-reviewed publication on wolf ecology in northwestern Manitoba (Parker 1973).

The overall purpose of this study was to employ a minimally invasive approach to examine spatial and temporal patterns of wolf occurrences within the summer range of the Eastern Migratory Cape Churchill caribou population in Manitoba, Canada as part of a larger comprehensive long-term caribou/wolf study. Our research objectives were informed by the traditional and local knowledge shared with us by the local people. We quantified the occurrences of wolves at each of three monitoring sites in the north, central, and south areas of the caribou summer range, in order to (1) determine any long-term change in wolf detection events across nine years of monitoring; (2) evaluate temporal and spatial patterns in seasonal wolf occurrences; and (3) characterize nocturnal and diurnal occurrences of wolves in each month. We predicted that (a) the annual frequency of wolf occurrences in this wilderness ecosystem would be relatively stable across years; (b) that wolf occurrences within caribou summer range would be approximately equal among the northern, central, and southern parts of the study area; (c) that most wolf occurrences would be during the summer months when caribou are present on their summer range and rare in winter when caribou are almost completely absent; and (d) that wolf occurrences would be highest during daylight hours and wolves would select strongly for daylight in winter when it is least available. These objectives and predictions were developed over the nine years of the study through engagement with local people who shared their knowledge and experience and through three community presentations in the town of Churchill. This is the first quantitative peer-reviewed ecological study to report on wolves in the Hudson Bay Lowlands of Manitoba.

## Methods

### Study area

Data were collected in Wapusk National Park (Wapusk;11,475 km^2^; Fig. 1), Manitoba, Canada in the Hudson Bay Lowlands of Manitoba within the Hudson Plains Ecozone (Wiken 1986). The study area boundary is based on the summer range of the Eastern Migratory Cape Churchill Caribou (Fig. 1), of which wolves are the top predator (Moayeri 2013). The study area is part of a broad transition zone between the lichen woodlands to the south and southwest and mostly treeless tundra to the north (Brook 2001). The study area is comprised of treeless poor fens (51%), sparsely vegetated gravel and sand beach ridges (16%), unvegetated shorelines and coastal ridges (13%), ponds, lakes, and rivers (8%), tall shrub fens (6%), sparsely treed fens and bogs (5%), and mostly degraded salt marsh (1%) (Brook 2001; Brook 2002). Summers in the study area are relatively short (June to August) and typically cool, while winters are long (November to March) and extremely cold, with consistently strong winds in all seasons.

**Fig. 1.**
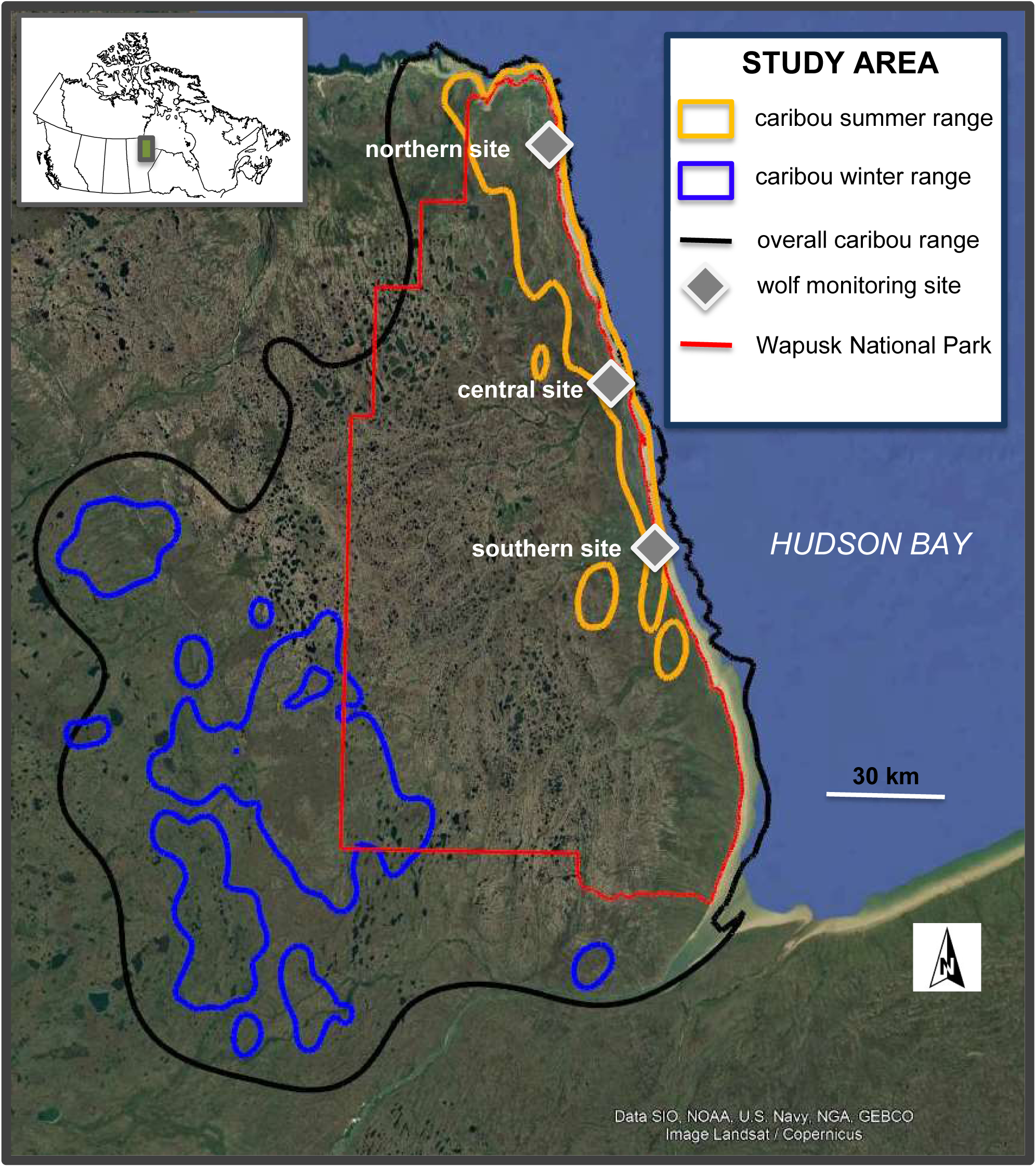
Map of the study area within the Greater Wapusk Ecosystem demonstrating the three monitoring sites within the summer range of the Eastern Migratory Cape Churchill caribou population.

Wolves in northern Manitoba have been hunted, trapped, and otherwise persecuted over the last century and a half by non-Indigenous people. Wolf bounties began in Manitoba in 1878 and mostly ended in 1965, in large part because they were overall ineffective (Pimlott 1961, Cluff and Murray 1995, Goldsborough 2008; Proulx and Rodtka 2015). Total annual payouts of wolf bounties peaked at $30,000 [equivalent to $660,810 in 2025 dollars] in 1917. In northern Manitoba an ostensible caribou ‘crisis’ emerged, based on hearsay and questionable science, starting in the 1920s (Usher 2004). This generated an intensive effort to reduce or eliminate wolves in migratory caribou range in northern Manitoba, initially with a bounty program that ended in the early 1950s (Usher 2004). A major wolf poisoning program was piloted in 1949 and implemented broadly until the early 1960s. Wolf populations took decades to recover after this (Usher 2004).

The previously unknown Hudson Bay subspecies of wolves (*C. lupus hudsonicus*) was first described by Goldman (1941) but was allied with the still contemporary *C. l. occidentalis* that exists to the west of the Hudson Bay subspecies. It was further recognized as a subspecies in Wozencraft (2005). NatureServe recognizes these wolves as ‘Secure’ (NatureServe Explorer 2025). In Manitoba, all wolves are legally classified as a game species so they can be hunted and trapped during current regulated seasons under the authority of any big game license, and baiting is allowed under certain restrictions for non-Indigenous people (Manitoba Government 2025).

Gray wolf hunting zone 2 surrounds Wapusk to the west and south. For non-Indigenous hunters who are Manitoba residents, the wolf season is open from August 21-March 31, and they are allowed one wolf per hunter each year. For non-Indigenous trappers, the study area includes both the Barrenlands District (6A) and Limestone District (6), and trappers may harvest wolves using firearms and traps; electronic calls can be used. Furthermore, wolves in Manitoba can legally be killed by property owners in defense of property without a trapping or hunting license. First Nations people with status are not required to have a hunting or trapping license, do not have restricted hours or seasons, have no bag limits, and are not subject to equipment restrictions in Manitoba and within Wapusk. Métis people in Manitoba have special harvesting rights but these do not currently extend into northern Manitoba (Manitoba Métis Federation 2024).

Dubois and Monson (2004) in an unpublished report summarized the state of knowledge of mammals in Wapusk and concluded that wolves were considered common within the Park. Other large predators in the study area included polar bears, black bears (*Ursus americanus*), and grizzly bears (*Ursus arctos)*, with polar bears being by far the most common (Clark et al. 2019). Wolverines (*Gulo gulo*) occurred sporadically (Dubois and Monson 2004, Clark unpublished data, 2024). Moayeri (2013) used stable isotope analysis to demonstrate that wolves in northern Manitoba had a high proportion of Cape Churchill caribou in their diet. Scurrah (2012) found that although the sample of satellite collared wolves in northeastern Manitoba was small (n=5), they move over very large areas. Wolves traveled 1,470 to 8,460 km per year and home ranges varied from 721 km^2^ to 196,020 km^2^ (Scurrah 2012). Historical work mapping 95 observations of wolf packs to the west and north of the study area in Manitoba and what is now Nunavut identified only 2 (2%) that were within 20 km of the Hudson Bay coast (Parker 1973).

### Field Data Collection

We deployed an average of five Reconyx PC-900 trail cameras at each of three survey locations that were 2-6 km inland from Hudson Bay (northern 58°N39’42.84”; 93°W11’26.52”, central 58°N7’50.16”; 92°W51’57.96”, and southern 57°N49’41.8794”; 92°W48’20.1594”; (Laforge et al 2017, Clark et al. 2019, Lochansky et al. 2025) within the summer range of the Eastern Migratory Cape Churchill Caribou (Fig. 1; Lochansky et al. 2025). The monitoring sites were based at three field camps that are used sparsely by researchers, university-based field courses, and park staff and can only be accessed by aircraft in summer and snowmobile or helicopter in winter.

Trail cameras were placed 85 cm above ground level and were bolted to heavy wooden poles buried 1.5m in the ground that formed a perimeter around each of the three field camps and were surrounded by heavy gauge paige wire to exclude polar bears (Laforge et al. 2017). Each camera was protected by a heavy duty metal Reconyx security enclosure. Trail cameras were serviced once per year to download all photos and replace the batteries. Trail cameras faced outwards with a minimum of one camera for each cardinal direction. The default trail camera settings (high detectability and high-resolution images) with a 46° field of view and 30m nominal detection range were used. The trail cameras were motion-activated and set to record three images at one second apart with a minimum 10s delay between activations. When triggered at night, the cameras used a ‘covert’ infrared flash which prevented disturbance of the animal (Gibeau and McTavish 2009). Trail cameras had 3.1 megapixel resolution and took HD color images in adequate ambient light levels and monochrome infrared images during nighttime.

All wolf photos were extracted yearly from each trail camera after the memory cards were downloaded and compiled into a single database, including survey location (northern, central, southern), date, time, year, and month. Images were coded individually into unique events that included all photos within a one hour window (Laforge et al. 2017, Clark et al. 2019). Six seasons were defined: early winter (November and December), mid-winter (January and February), late winter (March and April), spring (May and June), summer (July and August), and fall (September and October). Photos were labelled as taken at night when they were black and white and daytime when they were color images, based on the settings of the Reconyx cameras. Available daylight hours were determined using the National Research Council sunrise/sunset calculator and averaged for each month (Government of Canada 2024).

This research was approved by the University of Saskatchewan Animal Research Ethics Board, protocol number 20110076 and Parks Canada research permits WAP-2011-8336, WAP-2012-11956, WAP-2013-13933, and WAP-2017-25782.

### Data Analysis

We analyzed 418 unique wolf events documented over nine continuous years (2013-2021) of monitoring comprising 108 months (49,275 camera trap days) and compared the frequency of wolf occurrences at each site to a random distribution of occurrences using T-tests. Caribou were virtually absent from their summer range during the winter months. Overall, only 0.04% of caribou events were documented on their summer range during the winter snow period (November to April) since they migrate ∼150 km to their winter range to the southwest (Lochansky et al. 2025). Annual variation was characterized by conducting a regression analysis of the count of wolf events in each year over the nine years of the study. For any events that had > wolf photo, we calculated event length as the time between the first and last photo in each event. Seasonal variation was quantified by comparing wolf events in each of the previously defined six seasons and among the three monitoring sites. Temporal variation was assessed by analyzing monthly diurnal and nocturnal patterns of wolf occurrences. We compiled monthly temperature data collected by the trail camera images and reported the mean and standard error per month.

Selection Ratios (SRs) were calculated to determine if wolves were selecting or avoiding the nocturnal period of each month, as the ratio of the proportion used to the proportion available (Manly et al. 2002):

(w*_i_* = o*_i_*/π*_i_*) formula 1

where o*_i_* refers to the proportion of wolf occurrences at night, and π*_i_* represents the proportion of nocturnal time available. The selection threshold is 1.0. If use of nocturnal time is greater than it is proportionally available (i.e., selection of nocturnal time is occurring) the 95% confidence interval (CI) of the SR is >1. If the 95% CI of the SR is <1, nocturnal time is used less than available, (i.e., avoided); and if the 95% CI of the SR includes 1, nocturnal time is used as a function of its availability and is neither selected nor avoided.

## Results

We used trail camera data to describe wolf occurrence across days, seasons and years at a variety of light and temperature conditions in the summer range of the Eastern Migratory Cape Churchill Caribou in northern Manitoba, Canada. From 2013 to 2021 we collected 296, 144 photos from all trail cameras, of which 1.7% (n = 5,060) depicted wolves (see Fig. 2 for exemplary photos).

**Fig. 2.**
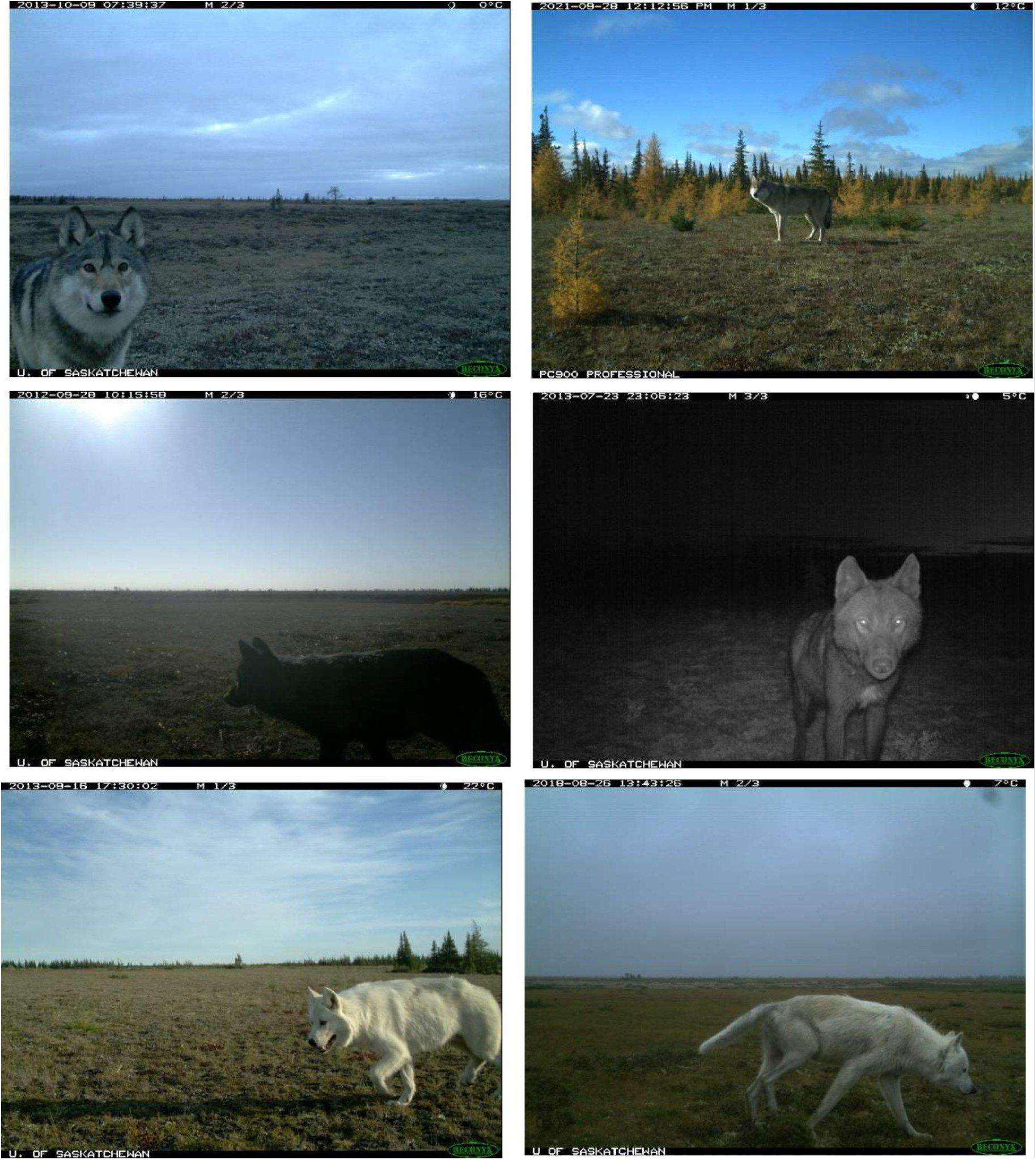
Examples of wolves detected by trail cameras at three monitoring sites along the Hudson Bay coast in Wapusk National Park, Manitoba, Canada (2013-2021).

We defined individual events as being separated by at least one hour and recorded 418 unique events that included wolves. Each event had an average of 11.5 photos per event (min = 1, max = 119). Of all of the wolf events, 8% were of a single photograph (i.e., event length = 0 seconds) and 73% were < 1 minute long. Of the events with >1 photograph, average event time was 2:59 minutes (min =0:01 minutes, max = 59:32 minutes).

Wolf events were documented at all monitoring sites in all years with one exception at the northern site in 2014. There was an average of 46.4 events per year over the nine years of the study (min = 20, max = 117) pooled across monitoring sites. The number of events did not change significantly across all years (R^2^= 0.055, df = 8, p = 0.515; Fig.3). Notably, 2017 was an outlier year with 117 events, more than 2.5x the average; the removal of the outlier did not change the trend in events across years (R^2^= 0.320, df = 7, p =0.113).

**Fig. 3.**
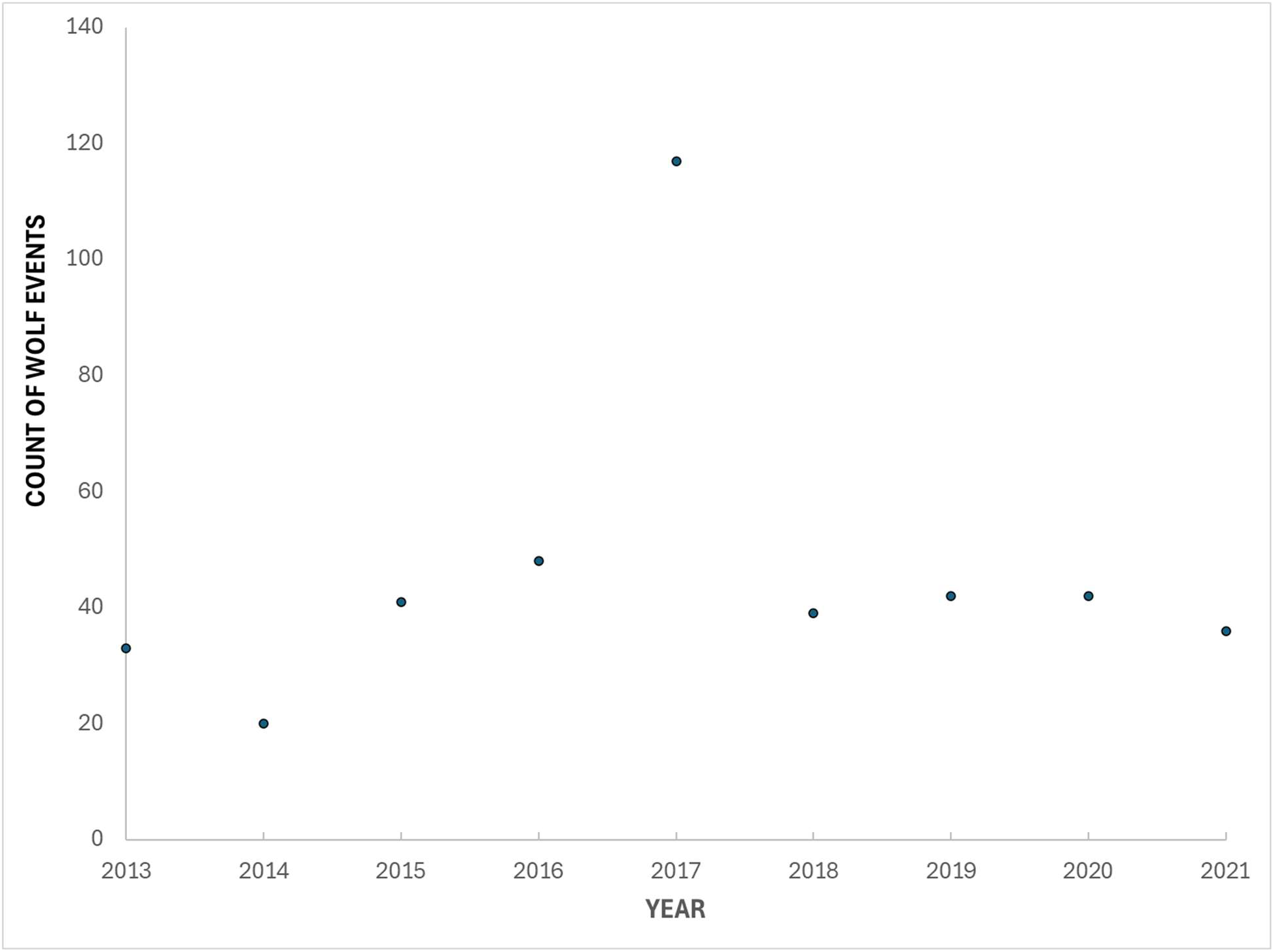
Annual trend in wolf events at trail cameras pooled across three monitoring sites by year along the Hudson Bay coast in Wapusk National Park, Manitoba, Canada (2013-2021; R=0.03, df=8, p <0.24) and this lack of significant trend remained when the outlier year 2017 was removed.

Wolves occurred at least once in 88 of the 108 months (81%). Annual counts of events at the southern monitoring site alone almost perfectly predicted the overall annual counts of all sites combined (R^2^= 0.990, df = 8, p < 0.0001). Across all nine years, 13 events included juveniles, all of which (100%) were at the southern monitoring site. Wolves also experienced extreme environmental conditions, with a 60°C range in temperature, from a low of −32°C in winter to a high of +28°C in summer and an annual change in day length of >11 hours between summer and winter (Fig.4).

**Fig. 4.**
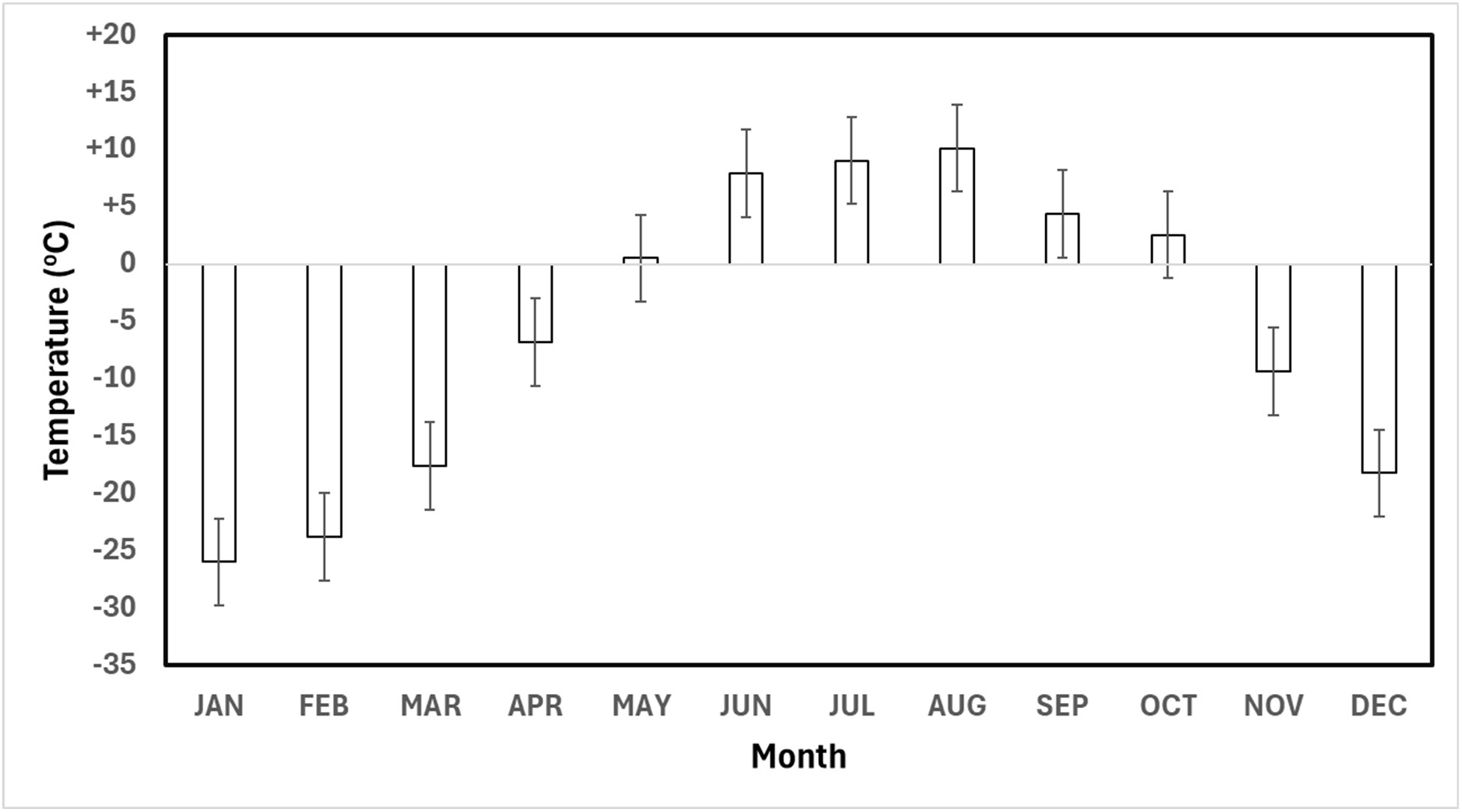
Average monthly temperature recorded by the Reconyx trail cameras on each photo image along the Hudson Bay coast in Wapusk National Park, Manitoba, Canada (2013-2021).

The average number of annual events at the Northern Site (mean =1.22 events/year) compared to a random distribution (15.5 occurrences per year per monitoring site) (mean =15.55 events/year) was significantly less than random, T= 39.192, df=8, p < 0.0001. The average number of annual events at the Central Site (mean =11.78 events/year) compared to random (mean =15.55 events/year) was significantly less than random T= 3.104, df=8, p = 0.0146. The average number of annual events at the Southern Site (mean =33.44 events/year) compared to random (mean =15.55 events/year) was not significantly different than random, T= −1.893, df=8, p = 0.0954, but when the 2017 outlier is removed, the Southern Site (mean =33.44 events/year) compared to random (mean =24.38 events/year) was significantly higher than random, T= −2.837, df=8, p < 0.0251, even though the monitoring areas were relatively close to each other (northern to central = 62 km, central to southern = 34 km).

Wolf events also varied significantly by season and by monitoring site (Fig. 5). Most (89%) wolf events occurred during the six month snow-free period (May to October) when caribou were absent; only 11% of wolf events occurred during the six month snow period (November to April) when caribou were absent (Fig. 5). Wolves were present in 100% of snow-free months across all years, but in only 43% of the months with snow. The year 2017 also had a larger proportion of events at the southern site; in all other years, the total events at the southern site averaged 61%, compared to 91% in 2017.

**Fig. 5.**
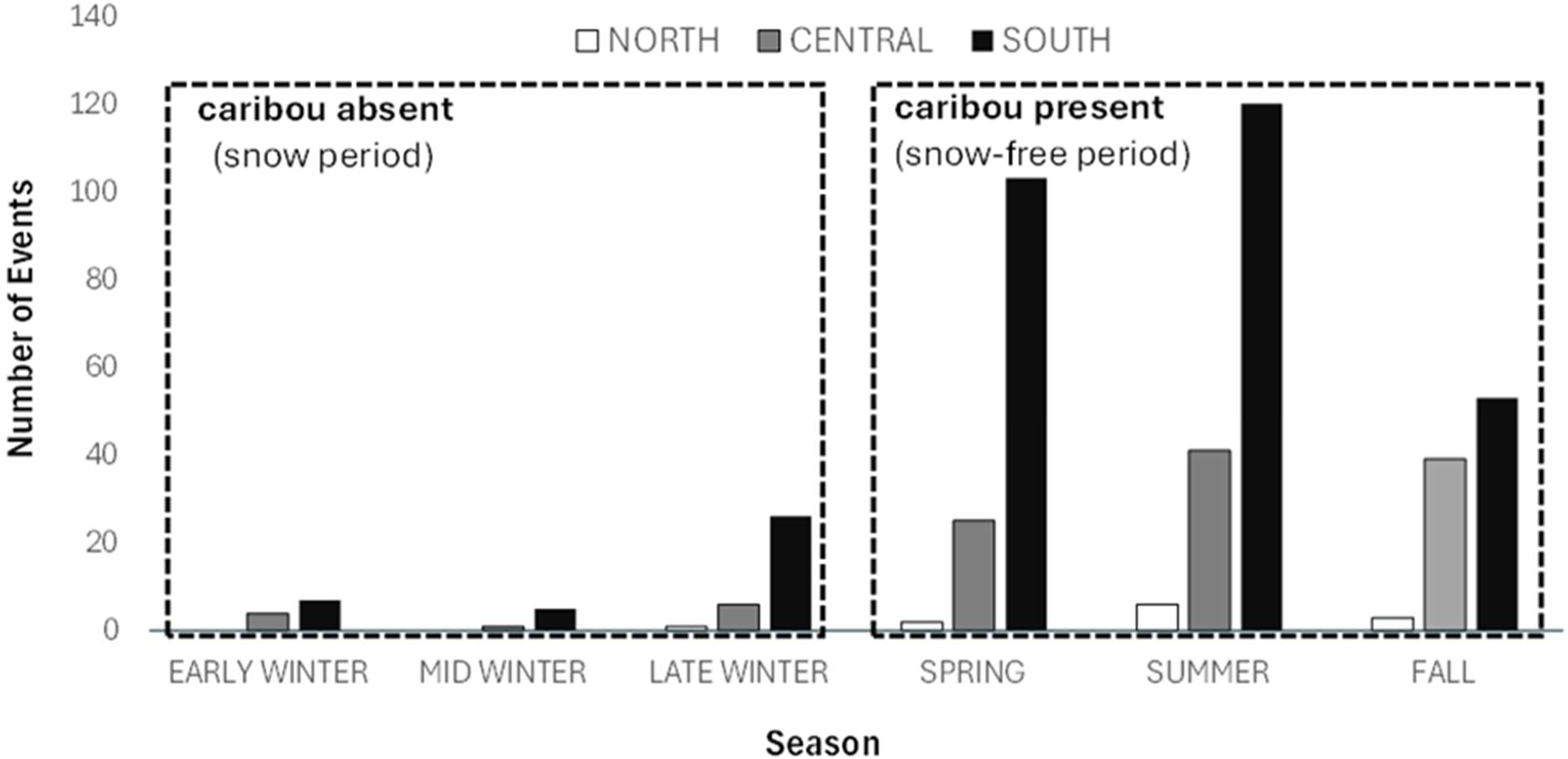
Seasonal patterns of events of wolves at field camps along the Hudson Bay coast in Wapusk National Park, Manitoba, Canada (2013-2021). Caribou presence in the study area is almost exclusively (99.9%) during the spring, summer, and fall seasons.

Within the study area, day length varied markedly over the 12 months of each year, from a low of 6 hours 14 minutes on December 21 to a high of 18 hours and 25 minutes on June 21, a difference of over 11 hours (Fig. 6). Of the 418 events detected across all years, 52% were during daylight hours and 48% were at night. This pattern varied monthly with nocturnal events being >50% of the total for the months of August, September, and October, though sample size was small in mid-winter (Fig. 6). Selection ratios (SR) indicated that wolves selected for being nocturnal in September (SR = 1.24), October (SR = 1.32), and November (SR = 1.29), though not strongly (i.e. >1.5) in any of these months (Fig. 7). Wolves were detected during nighttime less frequently than it was proportionally available to them (i.e., avoided night time) in January (SR = 0.00), February (SR = 0.53), March (SR = 0.60), April (SR = 0.74), May (SR = 0.52), June (SR = 0.42), and July (SR = 0.61; Fig. 7). Wolves neither avoided nor selected for nighttime in September (SR = 1.24) and December (SR = 1.00), using it at the same frequency at which it was available (Fig. 7).

**Fig. 6.**
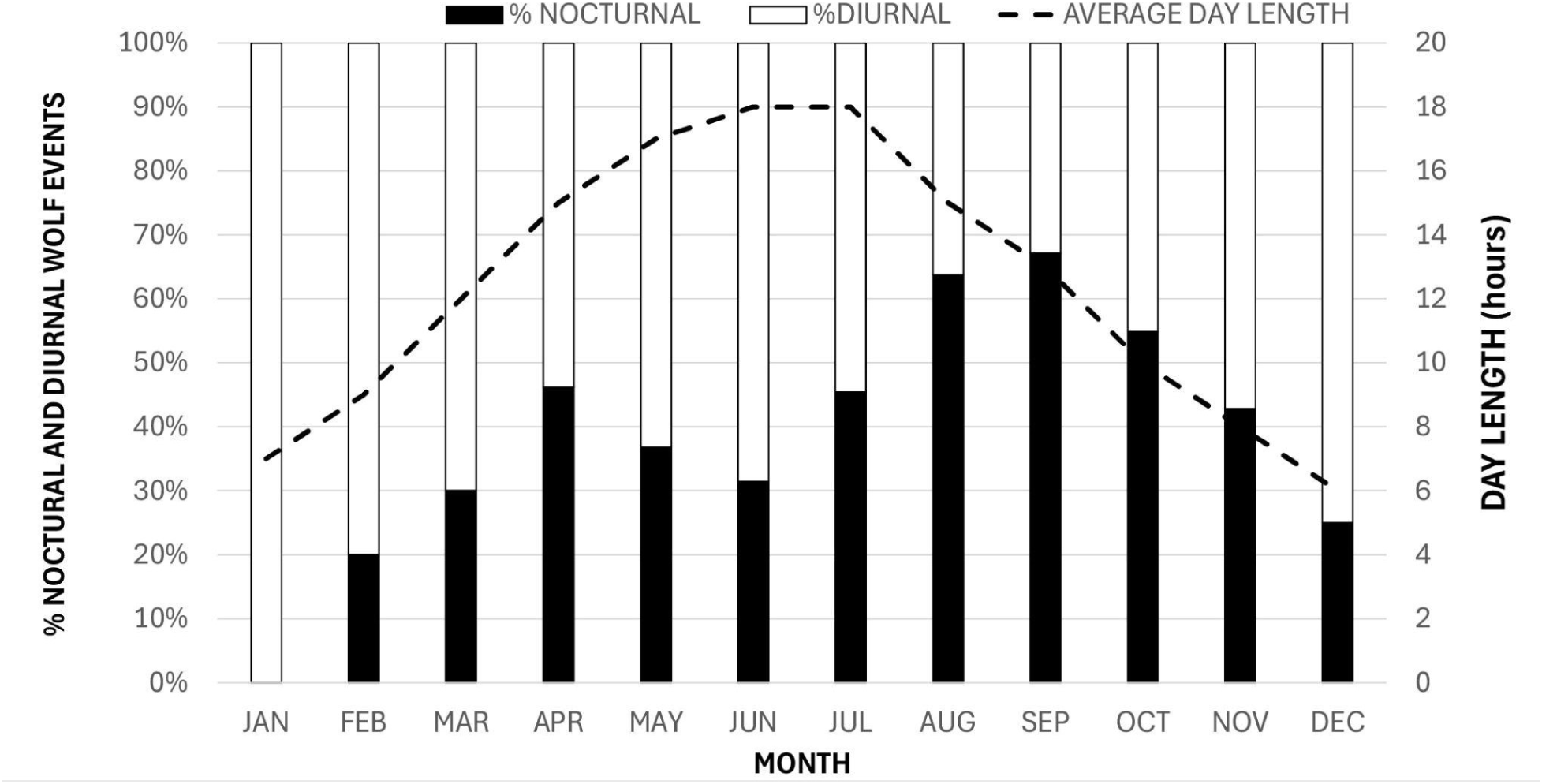
Monthly patterns in nocturnal and diurnal occurrences of wolves in the summer range of the Eastern Migratory Cape Churchill Caribou population along the Hudson Bay coast in Wapusk National Park, Manitoba, Canada (2013-2021).

**Fig. 7.**
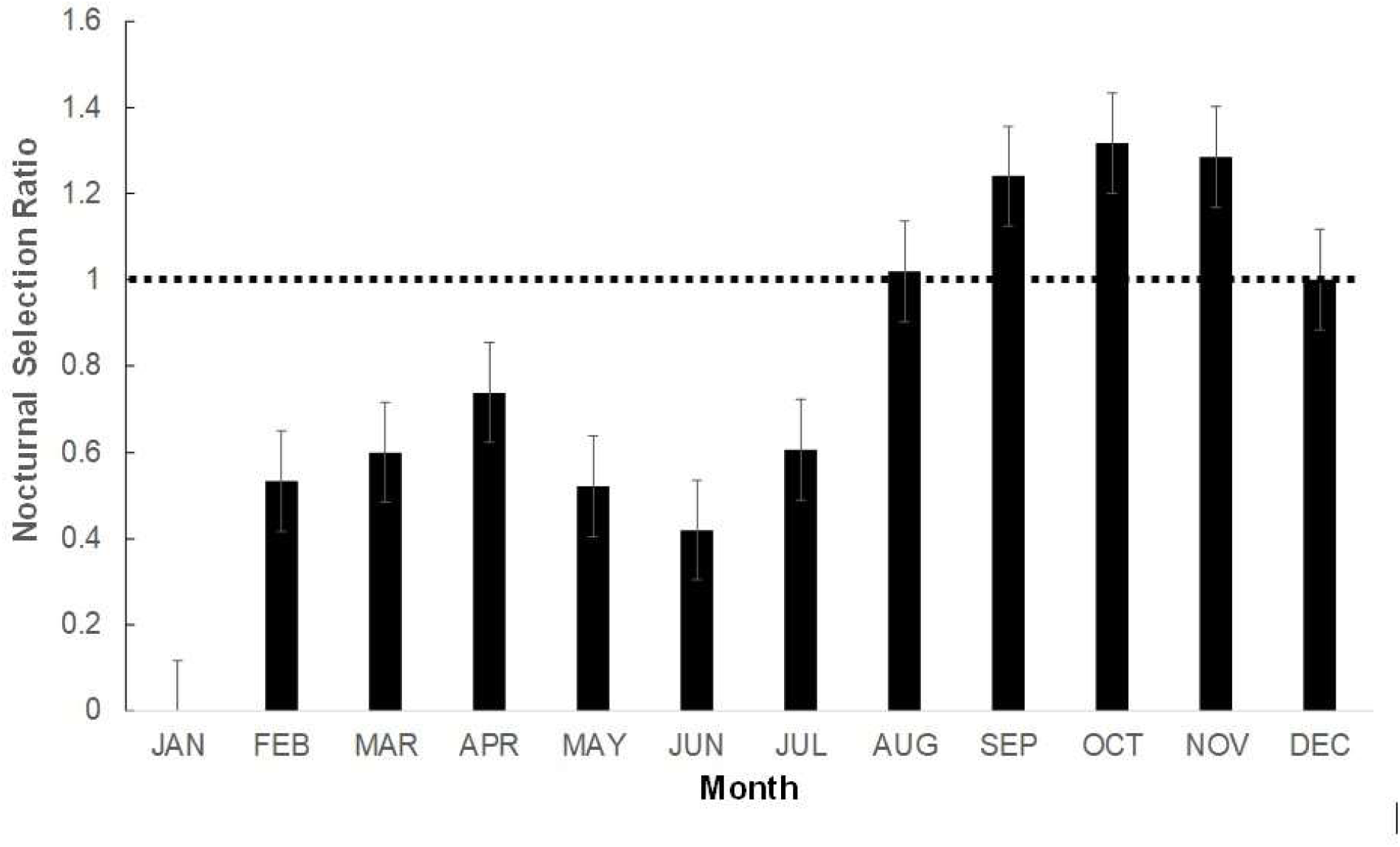
Wolf selection ratios for nighttime using selection ratios (% used / % available) in the summer range of the Eastern Migratory Cape Churchill Caribou population along the Hudson Bay coast in Wapusk National Park, Manitoba, Canada (2013-2021). Selection ratios below and not overlapping with 1 (e.g. January-July) indicate wolves use darkness less than it is available (i.e. avoid), SR above and not overlapping with 1 (e.g. September, October, and November) indicate wolves select darkness more than it is available. Any SR’s overlapping with 1 (e.g. August and December) indicated wolves neither selected or avoided darkness.

Trail cameras were deployed in clusters of 5 at each of the 3 monitoring sites and monitoring effort was consistent and continuous across the nine years of the study. All trail cameras operated continuously for the entire 9 year study period (2012-2021), though at times individual trail cameras failed for up to five weeks. However, >1 trail camera was always functioning at each monitoring site so there was no notable detection loss at any given location. The only notable issue was in 2020 when 2 trail cameras failed at the northern monitoring site due to internal clock battery failure, reducing monitoring effort for part of that year. Despite this, there were more wolves (n = 4 events/year) detected at the northern monitoring site in 2020 than in any other year of the study (max = 1 event/year; results not shown). Additionally, there were some periods in mid and late winter when individual trail cameras on the east side of camps were partially or completely covered in snow, but there was always >1 trail camera operating at each monitoring site and there is no evidence this affected overall detection rates. With trail cameras placed in each cardinal direction at each monitoring site, wolves were detected in all directions at all monitoring sites.

## Discussion

Our findings from this first ever quantitative peer-reviewed study of wolf occurrences in the Hudson Bay Lowlands in Manitoba determined that wolves were well established in Wapusk within the summer range of the Eastern Migratory Cape Churchill caribou population, supporting past unpublished work and local and traditional knowledge (Dubois and Monson 2004, Scurrah 2012). Consistent with our prediction, wolves were broadly distributed across our study area and were found consistently at all monitoring sites and in nearly all years. Contrary to our prediction that wolves would be randomly distributed, the most striking pattern overall was that the wolves were consistently concentrated in the most southern part of the study area and were relatively rare in the north. However, there are known wolf dens around the southern monitoring area that likely at least partly explains this pattern and the extremely high event year of 2017 is most likely when a den was active near the southern study site and the pack was closely associated with the den. Indeed, an even larger majority than normal of wolf events that year were at the southern site. While the overwhelming majority of caribou occurrences on these cameras was during the snow-free period (May to October), some caribou may have started using their summer range in late winter farther inland away from where the cameras were situated. This likely explains the increase in wolves detected on cameras during that period. Summer diet reconstruction from wolves in the study area by Moayeri (2013) found that the diet was, as expected, mostly caribou. While there is no available research on wolf winter diets, they are likely consuming more diverse prey in winter when caribou are absent, which may include moose (*Alces alces*), arctic hare (*Lepus arcticus*), snowshoe hare (*Lepus americanus*), rock ptarmigan (*Lagopus muta*), and willow ptarmigan (*Lagopus lagopus*). Furthermore, the exceptionally large home ranges of these wolves means that while visiting the southern monitoring site near the Hudson Bay coast, they likely also cover many different habitats, and they can readily move the 150 km between caribou summer and winter range over a short period of days and weeks.

Consistent with our prediction, wolf events occurred during daylight hours in >50% of all months except August, September and October and showed positive selection for nocturnal time in these same three months. Wolf use and selection for daylight occurred primarily when day length was increasing. Diurnal and nocturnal activity of wolves has been relatively unstudied in North America. Most of the available research has mainly focused on anthropogenic disturbances influencing wolf selection for nocturnal movements to avoid human conflict in Europe (Theuerkauf 2009; Pretiou et al. 2023; Ferreiro-Arias et al. 2024; Martínez-Abraín et al. 2023; Smith et al. 2024; Sunde et al. 2024). However, given the almost complete absence of human activity in Wapusk, especially at the southern monitoring site, wolves can be active during daylight without increased risk of disturbance or being killed by humans. Positive selection for the nocturnal period in later summer and fall is likely because caribou must increase hunting time considerably when caribou are spread out so much more widely across the summer range and are beginning to disperse out of summer range. Indeed, wolf diel activity positively correlates with increased prey availability (Sunde et al. 2024), which could explain wolf diurnal activity patterns in May-July when caribou first arrive in their summer range and day length is rapidly increasing. However, due to the lack of available research on wolf diel activity, we are unable to determine why wolves are selecting diurnal movement patterns for >50% of months of the year. Further research on diel patterns of caribou behaviour may help better understand this pattern.

Our study was limited by sample size, as we had only three monitoring sites in the study area, as well as some brief periodic camera failures. Future monitoring of wolves should involve deploying a more expansive network of trail cameras to capture the broader distribution of wolves in the region and cover both the summer range of the Eastern Migratory Cape Churchill caribou population and the winter range that also includes Qamanirjuaq and Penn Island migratory populations, as well as local bands of non-migratory caribou (Thomas and The Beverly and Qamanirjuaq Caribou Management Board 1996, Klütsch et al. 2016). Importantly, the caribou summer range is almost completely protected by Wapusk, while wolf trapping and hunting is legal in most of the caribou winter range. This area has limited protection in the Churchill Wildlife Management Area (CWMA) and no established habitat protections on all crown lands west of the CWMA where caribou and wolves are present during winter. While fires are absent from the mostly treeless summer range, they do occur extensively throughout the largely treed winter range, and this may also influence both caribou and wolf movements. Next steps should include a detailed comparison between wolf and caribou occurrences and seasonal habitat selection across both summer and winter caribou ranges and at individual cameras.

Additional analysis should include factors influencing wolf activity, including group size, moon phase, and habitat. Studies of wolves outside of the protections and remoteness provided by Wapusk are necessary for our understanding of wolf ecology in northern, remote ecosystems.

As northern communities expect and increasingly demand less invasive research and management and often call for greater involvement in wildlife research and management, trail camera research creates important opportunities to engage local communities in deploying and monitoring their own cameras (Kemp et al. 2024). Additionally, the potential for engaging youth in research and monitoring is an important opportunity for community engagement (Brook et al. 2009). Research linking local and traditional knowledge with trail camera data represents a significant opportunity to obtain vital long-term data and engage communities in meaningful ways (Brook and McLachlan 2005). Additional trail cameras to monitor wolves outside of Wapusk where they are trapped, shot, and at times disturbed by humans would be prudent.

From a practical perspective, there are no direct management options for wolves within a remote wilderness National Park in Canada, such as Wapusk where the management board is strongly committed to avoiding any human interventions and overall use of Wapusk by local people is low (Government of Canada 2017). While trapping of other fur-bearing animals and hunting of caribou is allowed under specific conditions, there is no legal trapping or shooting of wolves in Wapusk by non-Indigenous people (Government of Canada 2010). While treaty rights to harvest are recognized, currently in practice, Indigenous trapping and hunting of wolves in this area is nearly non-existent(Government of Canada 2017). However, the research presented here and ongoing monitoring using trail cameras currently serves as the only quantitative indicator of wolf population patterns and is crucial for the early detection of future population decline in the region.

We recommend that if less than 29 wolf events (average - one standard deviation) are detected annually for at least three years, that should be considered a potential concern. Below average wolf occurrences should trigger additional research on wolf ecology, parasites, disease monitoring, and climate effects. While low wolf populations are unlikely to trigger any direct management responses within Wapusk, they should be used to consider adjusting wolf trapping and hunting opportunities outside of Wapusk. We do not expect that the trail camera data alone would ever indicate an ‘over-abundance’ of wolves, or that there would (or should) be a management response in that regard. However, if annual wolf events are greater than 46 wolf events annually (average + one standard deviation) for more than three years, that should trigger an evaluation or consideration of trapping and hunting in the area around Wapusk, but only if there is also compelling evidence that prey species such as caribou are significantly declining. At the same time, research and monitoring is needed on the primary prey source of the wolf population, the Eastern Migratory Cape Churchill caribou, and other overlapping caribou populations on the caribou winter range to determine cow:calf ratios and estimate population sizes annually. Our findings that wolf occurrences appear stable in the caribou summer range, and that wolves are consistently identified annually at northern, central, and southern monitoring sites each year is encouraging that populations have not been critically low during the time we have been monitoring them.

## Acknowledgements

First, we thank Dr. Paul Paquet who is the ultimate wolf expert and mentor. We also recognize Farley Mowat who inspired our research on wolves on the Hudson Bay coast with ‘Never Cry Wolf’. We thank Gordy Kidlapik and Florence Hamilton for sharing Indigenous names for wolves. We gratefully acknowledge field support from Hudson Bay Helicopters and numerous excellent and safe pilots, the Churchill Northern Studies Centre, Wapusk National Park staff, and numerous individuals including Murray Gillespie, Sheldon Kowalchuk, Melissa Gibbons, Jill Larkin, Jessica Lankshear, Natalie Asselin, Heather MacLeod, David Britten, Nils Lokken, Aimee Schmidt, James Roth, Frank Baldwin, James Baldwin, Julie Rogers, Matt Webb, Kassandra Peterson, and Brianne Symak.

## Data Availability

Data generated or analyzed during this study are subject to an embargo of 24 months from the publication data of this manuscript. Once the embargo expires, the data will be available upon reasonable request to the corresponding author.

## Author contributions

Conceptualization: RKB, DAC Data curation: RKB, DAC Formal analysis: RKB

Funding acquisition: DAC, RKB Investigation: RKB, KH, DAC, CL Methodology: RKB

Project Administration: RKB, DAC Resources: RKB, DAC Supervision: RKB, DAC

Writing-original draft: RKB

Writing-review & editing: RKB, KH, DAC, CL, JC

---

**Table 1.**
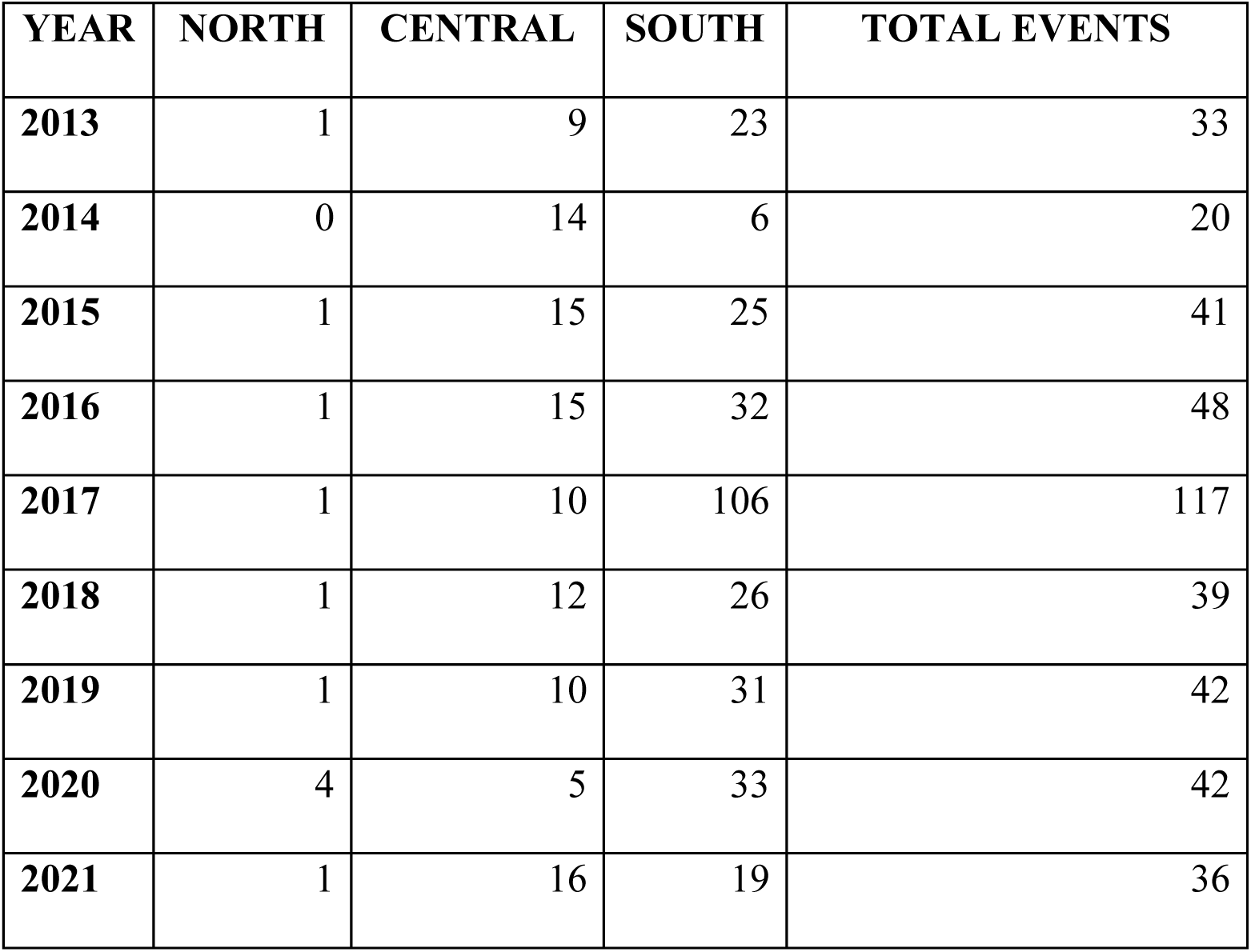
Spatial and temporal summary of wolf events detected in the Reconyx trail cameras along the Hudson Bay coast in Wapusk National Park, Manitoba, Canada (2013-2021).

## References

Ausband, D. E., Lukacs, P. M., Hurley, M., Roberts, S., Strickfaden, K., and A.K., Moeller. 2022. Estimating wolf abundance from cameras. Ecos., 13(2). 10.1002/ecs2.3933

Anderson, M.L., Cluff, H.D., Mech, L.D., and MacNulty, D.R. 2024.Wolf density and predation patterns in the Canadian High Arctic. J. Wildl. Manage. 89:e22671. 10.1002/jwmg.22671

Barber-Meyer, S.M. 2022. Can non-invasive methods replace radio collar-based winter counts in a 50-year wolf study? Lessons learned from a three-winter trial. Wildl. Res. 50(6):451–464. 10.1071/WR22001

Barclay, P.D. 2010. A ’Curious and Grim Testimony to a Persistent Human Blindness’: Wolf Bounties in North America, 1630-1752. Eth. Plac. Envir. 5(1):25–34. 10.1080/13668790220146429

Blossey, B., and Hare, H. 2022. Myths, Wishful Thinking, and Accountability in Predator Conservation and Management in the United States. Front. Cons. Sci. 3. 10.3389/fcosc.2022.881483

Boiseau, B.T., Trinidad, J.M., Knight, R.N., Larsen, R.T., McMillan, B.R., and Hall, L.K. 2024. Influence of moonlight on visits to water sources by mammalian predator and prey: a test of competing hypotheses. Animal Behaviour 210: 139–152. 10.1016/j.anbehav.2024.02.005

Bonin, M., Dussault, C., Taillon, J., Pisapio, J., Lecomte, N., and Côté, S. 2023. Diet flexibility of wolves and black bears in the range of migratory caribou. Journal of Mammalogy 104. 10.1093/jmammal/gyad002

Botts, R.T., Eppert, A.A., Wiegman, T.J., Blankenship, S.R., Rodriguez, A., Wagner, A.P., Ullrich, S.E., Allen, G.R., Garley, W.M., Asselin, E.M., and Mooring, M.S. 2020. Does Moonlight Increase Predation Risk for Elusive Mammals in Costa Rica? Tropical Conservation Science 13: 1940082920952405. 10.1177/19400829209524

Brook, R.K. 2001. Structure and dynamics of the vegetation in Wapusk National Park and the Cape Churchill Wildlife Management Area of Manitoba, community and landscape scales. Master of Natural Resource Management, University of Manitoba. http://hdl.handle.net/1993/2724

Brook, R. K., and Kenkel, N. C. 2002. A multivariate approach to vegetation mapping of Manitoba’s Hudson Bay Lowlands. Intern. J. Rem. Sens. 23(21), 4761–4776. 10.1080/01431160110113917

Brook, R.K., Cattet, M., Darimont, C.T., Paquet, P.C., and G.Prouix. 2015. Maintaining Ethical Standards During Conservation Crises. Can. Wildl. Biol. Manag., 4:72–79

Chavez, A.S. and Gese, E.M. 2006. Landscape use and movements of wolves in relation to livestock in a wildland-agriculture matrix. J. Wild. Manag., 70(4), pp.1079–1086. 10.2193/0022-541X(2006)70[1079:LUAMOW]2.0.CO;2

Clare, J.D. J., Zuckerberg, B., Liu, N., Stenglein, J. L., Van Deelen, T. R., Pauli, J. N., AND P.A. Townsend. 2023. A phenology of fear: Investigating scale and seasonality in predator–prey games between wolves and white-tailed deer. Ecol., 104(5). 10.1002/ecy.4019

Clark, D.A., Brook, R.K., Oliphant-Reskanski, C., Laforge, M.P., Olson, K., and Rivet, R. 2019. Novel range overlap of three ursids in the Canadian subarctic. Arc. Sci. 5: 62–70 (2019) dx.doi.org/10.1139/as-2018-0013

Cluff, H.D and Murray, D.L. Review of wolf control methods in North America. In Ecology and Conservation of Wolves in a Changing World; Carbyn, L.N., Fritts, S.H., Seip, D.R., Eds.; Canadian Circumpolar Institute, University of Alberta: Edmonton, AB, Canada, 1995; pp. 491–504

Creel, S., Schuette, P., and Christianson, D. 2014. Effects of predation risk on group size, vigilance, and foraging behavior in an African ungulate community. Behav. Ecol. 25(4): 773–784. 10.1093/beheco/aru050

Dalerum, F., Freire, S., Angerbjörn, A., Lecomte, N., Lindgren, A., Meijer, T., Pečnerová, P., and Dalén, L. 2018. Exploring the diet of arctic wolves (*Canis lupus arctos*) at their northern range limit. Canadian Journal of Zoology. 96(3): 277–281. 10.1139/cjz-2017-0054

David, A., Latham, M., Latham, C., Boyce, M.S., Boutin, S. 2011. Movement responses by wolves to industrial linear features and their effect on woodland caribou in northeastern Alberta. Ecological Applications 21: 2854–2865. https://doi.org/10.1890/11-0666.1

Dubois, J., Monson, K., 2004. Mammals of Wapusk National Park: survey results and a provisional checklist. Blue Jay 62 (3): 160–166. 10.29173/bluejay5993

Fancy, S.G. and Ballard, W.B. 1995. Monitoring wolf activity by satellite. Ecology and conservation of wolves in a changing world 35: 329–333. Monitoring Wolf Activity bySatellite

Ferreiro-Arias, I., García, E.J., Palacios, V., Sazatornil, V., Rodríguez, A., López-Bao, J.V. and Llaneza, L., 2024. Drivers of Wolf Activity in a Human-Dominated Landscape and Its Individual Variability Toward Anthropogenic Disturbance. Ecol. Evol. 14(10), p.e70397. 10.1002/ece3.70397

Frame, P.F., Hik, D.S., Cluff, H.D., and Paquet, P.C. 2004. Long foraging movement of a denning tundra wolf. Arctic 57: 196–203. https://www.jstor.org/stable/40512619

Goldsborough, G., 2008. Manitoba’s war on wildlife. Man. Hist. (59), pp.40-47.

Government of Canada. 2010. Wapusk National Park of Canada Park Use Regulations. SOR/2010-67. [accessed April 7, 2025]. https://laws-lois.justice.gc.ca/eng/regulations/sor-2010-67/FullText.html

Government of Canada. 2017. Wapusk National Park of Canada Management Plan. [accessed April 7, 2025] https://parks.canada.ca/pn-np/mb/wapusk/info/gestion-management/gestion-management-2017

Gibeau, M.L., McTavish, C. 2009. Not-so-candid cameras: how to prevent camera traps from skewing animal behaviour. Wildl. Profess. [accessed April 7, 2025] Not_so_candid_cameras.pdf

Goldman, E.A. 1941. Three new Wolves from North America, Proc. Biol. Soc. Wash. 54: 109–113.

Goldsborough, G. Manitoba’s War on Wildlife. https://www.mhs.mb.ca/docs/mb_history/59/waronwildlife.shtml

Government of Canada. 2017. Wapusk National Park Management Plan. [accessed on november 22, 2024]. Wapusk National Park Management Plan, 2017 - Wapusk National Park

Government of Canada. 2024. Caribou Conservation. [accessed on November 22, 2024]. Caribouconservation - Wapusk National Park

Hansen, J. and C. Urbigkit. 2021. Monitoring for Wolves. Wildlife Damage Management Technical Series. USDA, APHIS, WS National Wildlife Research Center. Fort Collins, Colorado. 11p. “Monitoring for Wolves” by Jeff Hansen and Cat Urbigkit

Johnson, C. J., Ray, J. C., & St-Laurent, M. H. 2022. Efficacy and ethics of intensive predator management to save endangered caribou. Cons. Sci. Prac. 4(7), e12729. 10.1111/csp2.12729

Klütsch. C.F.C., Manseau, M., Trim, V., Polfus, J., and Wilson, P.J. 2016. The eastern migratory caribou: the role of genetic introgression in ecotype evolution. Roy. Soc. Op. Sci. 3: 150469. 10.1098/rsos.150469

Latham, A.D.M., Latham, M.C., Boyce, M.S. and S. Boutin. 2011. Movement responses by wolves to industrial linear features and their effect on woodland caribou in northeastern Alberta. Ecol. Appl., 21(8): 2854–2865. 10.1890/11-0666.1

Laforge, M., Clark, D., Schmidt, A., Lankshear, J., Kowlachuk, S., and Brook, R. 2017. Temporal aspects of polar bear (*Ursus maritimus*) occurrences at field camps in Wapusk National Park, Canada. Pol. Biol. 40: 1661–1670. doi:10.1007/s00300-017-2091-6

Lochansky, C., Boudreau, M.R., Turner, R., Kliewer, M., Webb, M., Clark, D. and Brook, R.K., 2025. Temperature drives summer group size dynamics of Eastern Migratory caribou in the Hudson Bay lowlands of Manitoba. Can. J. Zool., 103, pp.1–8. 10.1139/cjz-2024-0062

Walton, L.R., Cluff, H.D., Paquet, P.C., and Ramsay, M.A. 2001. Movement Patterns of Barren-Ground Wolves in the Central Canadian Arctic, J. Mammal., 82(3): 867–876. https://www.jstor.org/stable/1383622

Manitoba Municipal Act, 1873. Winnipeg, Manitoba, Canada. [accessed April 7, 2025] Manitoba Laws

Manitoba Métis Federation. 2024. The Métis of the Harvest, revised 3rd edition. https://www.mmf.mb.ca/wcm-docs/docs/harvesters/me_769_tis_laws_of_the_harvest_2024.pdf

Manly, B. F. J., McDonald, L., Thomas, D., McDonald, T. L., & Erickson, W. P. (2002). Resource selection by animals (2nd ed.). Dordrecht, Netherlands: Kluwer Academic Publishers.

Martínez-Abraín, A., Llinares, Á., Llaneza, L., Santidrián Tomillo, P., Pita-Romero, J., Valle-García, R.J., Formoso-Freire, V., Perina, A. and Oro, D. 2023. Increased grey wolf diurnality in southern Europe under human-restricted conditions. J. Mammal. 104(4):846–854. doi: 10.1093/jmammal/gyad003

Michelot, C., Leclerc, M., Taillon, J., Dussault, C., Hénault Richard, J., and Côté, S.D. 2024. Evidence of migratory coupling between grey wolves and migratory caribou. Oikos e10150. 10.1111/oik.10150

Mishra, S. 2024. Keystone species and their impact on the ecosystem: Curr. Trend. Seed., 12: 13. doi10.61577/jec.2024.100001

Moayeri, M. 2013. Reconstructing the Summer Diet of Wolves in a Complex Multi-Ungulate System in Northern Manitoba, Canada. Master of Science Thesis, University of Manitoba. content

Musiani, M., Leonard, J.A., Cluff, H.D., Gates, C.C., Mariani, S., Paquet, P.C., Vilà, C., and Wayne, R.K. 2007. Differentiation of tundra/taiga and boreal coniferous forest wolves: genetics, coat colour and association with migratory caribou. Mol. Ecol. 16(19): 4149–4170. 10.1111/j.1365-294X.2007.03458.x

Packard, M. Jane. 2006. Wolf Behavior: Reproductive, Social, And Intelligent. In Wolves: behavior, ecology, and conservation, Repr. Edited by L.D. Mech and L. Boitani. Univ. of Chicago Press, Chicago, Ill.

Packard, J., Gordon, W. and J. Clarkson. 2011. Chapter 5. Biodiversity. In The Impact of Global Warming on Texas: Second edition. University of Texa Press: 124–156. doi.org/10.7560/723306-009

Parker, G.R. 1973. Distribution and densities of wolves within barren-ground caribou range in northern mainland Canada. J. Mammal. 54: 341–348. https://doi.org/10.2307/1379121

Petridou, M., Benson, J. F., Gimenez, O. and Kati, V. 2023. Spatiotemporal patterns of wolves, and sympatric predators and prey relative to human disturbance in northwestern Greece. Divers. 15:18.

Pimlott, D.H. 1961. Wolf control in Canada. Can. Audubon, 23, 145–152.

Proulx, G. and Rodtka, D., 2015. Predator bounties in Western Canada cause animal suffering and compromise wildlife conservation efforts. Animals, 5(4), pp.1034–1046. 10.3390/ani5040397

Proulx, G. and Parr, S. 2018. Is Livestock an Important Food Resource for Coyotes and Wolves in Central Eastern Alberta Counties with Predator Control Bounties? Can. Wild. Biol. Manag., 7(1):31–45.

Rafiq, K., Jordan, N.R., Golabek, K., McNutt, J.W., Wilson, A., and Abrahms, B. 2023. Increasing ambient temperatures trigger shifts in activity patterns and temporal partitioning in a large carnivore guild. Proc. Roy. Soc. B. Biol. Sci., 290(2010): 20231938.

Ramsay, M.A., Stirling, I. 1984. Interactions of wolves and polar bears in Northern Manitoba. J. Mammal., 65(4):693–694. 10.2307/1380856

Rolf, P., O., and Paolo, C. 2006. The Wolf as a Carnivore. In Wolves: behavior, ecology, and conservation, Repr. Edited by L.D. Mech and L. Boitani. Univ. of Chicago Press, Chicago, Ill.

Rossa, M., Lovari, S. and Ferretti, F., 2021. Spatiotemporal patterns of wolf, mesocarnivores and prey in a Mediterranean area. Behav. Ecol. Sociobiol., 75: 1–13. 10.1007/s00265-020-02956-4

Schmitz, O. J., D. Hawlena, and G. R. Trussell. 2010. Predator control of ecosystem nutrient dynamics. Ecol. Lett., 13: 1199–1209. 10.1111/j.1461-0248.2010.01511.x

Scurrah, F.E., 2013. Gray wolves (Canis lupus) movement patterns in Manitoba: implications for wolf management plans. Royal Roads University. DOI: 10.13140/RG.2.2.32003.9680

Smith, A.F., Kasper, K., Lazzeri, L., Schulte, M., Kudrenko, S., Say-Sallaz, E., Churski, M., Shamovich, D., Obrizan, S., Domashevsky, S. and Korepanova, K. 2024. Reduced human disturbance increases diurnal activity in wolves, but not Eurasian lynx. Glob. Ecol. Cons., p.e02985. 10.1016/j.gecco.2024.e02985

Sunde, P., Kjeldgaard, S.A., Mortensen, R.M. and Olsen, K. 2024. Human avoidance, selection for darkness and prey activity explain wolf diel activity in a highly cultivated landscape. Wildl. Biol., p.e01251. 10.1002/wlb3.01251

Theuerkauf, J., Jedrzejewski, W., Schmidt, K., Okarma, H., Ruczyński, I., Śniezko, S., and Gula, R. 2003. Daily Patterns and Duration of Wolf Activity in the Białowieza Forest, Poland. J. Mammal., 84: 243–253. doi:10.1644/15451542(2003)084%3C0243:DPADOW%3E2.0.CO;2.

Theuerkauf, J. 2009. What Drives Wolves: Fear or Hunger? Humans, Diet, Climate and Wolf Activity Patterns. Ethol., 115(7): 649–657. 10.1111/j.1439-0310.2009.01653.x

Thomas, D.C., and The Beverly and Qamanirjuaq Caribou Management Board. 1996. A fire suppression model for forested range of the Beverly and Qamanirjuaq herds of caribou. Rangif., 9:343–350. aysaekanger,+1276-4871-1-CE.pdf

Ripple, W.J., Beschta, Wolf, C., Painter, L.E., and Wirsing, A.J. 2025. The strength of the Yellowstone trophic cascade after wolf reintroduction. Glob. Ecol. Cons., 58: e03428. 10.1016/j.gecco.2025.e03428 https://doi.org/10.1016/j.gecco.2025.e03428

Usher, P. J. Caribou Crisis or Administrative Crisis? Wildlife and Aboriginal Policies on the Barren Grounds of Canada, 1947–60, in Cultivating Arctic Landscapes: Knowing and Managing Animals in the Circumpolar North, D.G.Anderson,M.Nuttall, Eds. (Berghahn Books, 2004), pp. 172–199. doi.org/10.1515/9781782382096-015

Vilà, C., Urios, V., and Castroviejo, J. 1995. Observations on the daily activity patterns in the Iberian wolf. [accessed April 7, 2025] https://www.semanticscholar.org/paper/Observations-on-the-daily-activity-patterns-in-the-Vil%C3%A0-Urios/c3fb85e967b1f65793f25ed76db17c409945b1b0

Walton, L.R., Cluff, H.D., Paquet, P.C.,Ramsay, M.A. 2001. Movement patterns of barren-ground wolves in the Central Canadian Arctic., J. Mammal. 82 (3): 867–876. 10.1644/1545-1542(2001)082<0867:MPOBGW>2.0.CO;2

Wiken, E.B. 1986. Terrestrial ecozones of Canada. Ecological Land Classification, Series No. 19. Environment Canada. Hull, Quebec. 26pp. + map. ia800804.us.archive.org/19/items/31761115540007/31761115540007.pdf

Winnie, J., and Creel, S. 2017. The many effects of carnivores on their prey and their implications for trophic cascades, and ecosystem structure and function. Food. Web. 12: 88–94. 10.1016/j.fooweb.2016.09.002

Wolfe, R. J. 2006. Playing with fish and other lessons from the North. University of Arizona Press.

Wozencraft, W. C. 2005. Order Carnivora. Pp. 512–628, in Mammal species of the world. A taxonomic and geographic reference (Wilson, D. E., and D. A. M. Reeder, eds.). Third edition. The Johns Hopkins University Press. Baltimore, U.S.A. https://doi.org/10.1644/06-MAMM-R-422.1

